# Differences in metagenome coverage may confound abundance-based and diversity conclusions

**DOI:** 10.1101/2024.10.10.617679

**Authors:** Borja Aldeguer-Riquelme, Luis M. Rodriguez-R, Konstantinos T. Konstantinidis

## Abstract

Although the importance of rarefying 16S rRNA gene data to the same coverage for reliable comparisons of diversity between samples has been well appreciated, the impact of (shotgun) metagenome coverage (i.e., what fraction of diversity was sequenced) on biological conclusions is commonly overlooked. We demonstrate that uneven coverage can lead to misleading conclusions about which features (e.g., genes, genomes) may differentiate two metagenomes or their diversity comparisons. We outline an approach to minimize this impact.

## Introduction

Rarefaction analysis attempts to account for (or normalize) differences in coverage between samples. Coverage here refers to the amount of sample’s total diversity recovered by the sequencing effort applied. Rarefaction has been shown to be necessary when comparing 16S rRNA gene amplicon datasets in order to identify differentially abundant taxa, differences in diversity and more between the datasets (1). Moreover, it is now well recognized that rarefying amplicons by sequencing effort is an insufficient, and often counter-productive method and coverage normalization should instead guide rarefaction (2). Despite the recognized importance of the latter metric, however, coverage is still largely ignored in comparative shotgun metagenomic studies. This is presumably because it is technically much more challenging to rarefy sequences that are not overlapping like in shotgun metagenomic datasets (or simply metagenomes) relative to overlapping amplicon data. Rodriguez-R and Konstantinidis developed Nonpareil in 2014 to calculate metagenome coverage based on read redundancy, which was termed Npc (3), and showed that the level of coverage has a notable effect on gene content comparisons between metagenomes (4). We further highlight the importance of metagenome coverage here by showing that overlooking coverage differences has led several published studies to inaccurate conclusions about what features (e.g., genes, pathways, or genomes) differ in abundance and/or diversity between microbiomes. We expand on our previous Nonpareil work by identifying the underlying cause for this limitation and proposing a simple approach to sidestep it.

## Results and discussion

We showcase below the impact of differences in Npc on derived conclusions by comparing the relative abundances of 219 metagenome-assembled genomes (MAGs) recovered from a marine metagenomic depth profile dataset published by Hawley and colleagues (5). These abundances were robustly estimated as Sequencing Depth normalized by Genome Equivalents (SD/GEQ, see Methods) in the original (full-size) metagenomes as well as in subsampled metagenomes of varying coverage levels, ranging from 0.1 to 0.7 Npc. The relative abundances of MAGs belonging to the same taxon (e.g., same order or family) were added up to represent the abundance of the taxon in each sample and the derived taxon abundances were directly compared. We consistently observed, across multiple taxa, an increase in their relative abundance in subsampled metagenomes of increasing Npc. For instance, the aggregated relative abundance of MAGs belonging to the phylum *Pseudomonadota* increased by about 26% in metagenomes of Npc=0.2 vs. 0.7, while that of *Thermoproteota* changed by (only) ∼2% in the same metagenome (**Figure 1A**). This result was counterintuitive since relative abundance should (theoretically) remain stable in subsets of the total metagenome given the presumed randomness and unbiased nature of shotgun metagenome sequencing (but see also (6, 7)). We observed that while the relative abundance of detectable individual MAGs remained stable in the subsampled metagenomes (**Figure 1C**), consistent with the above expectations, the number of MAGs detected increased progressively with increasing Npc, especially for *Pseudomonadota* MAGs (**Figure 1B**). These results indicated that sensitivity in detecting members of the feature, in this case *Pseudomonadota* and *Thermoproteota* MAGs, was responsible for the differences in total relative abundance of the feature observed as a function of the Npc (**Figure 1A**). In other words, the higher the sequencing effort applied (or Npc coverage), the higher the limit of detection and thus, the number of detected MAGs, which translates to higher aggregated abundance of the corresponding feature that the MAGs are assigned to. This effect was more pronounced for the *Pseudomonadota* than the *Thermoproteota* due to the higher number of low-abundance MAGs assigned to the former phylum.

**Figure 1.**
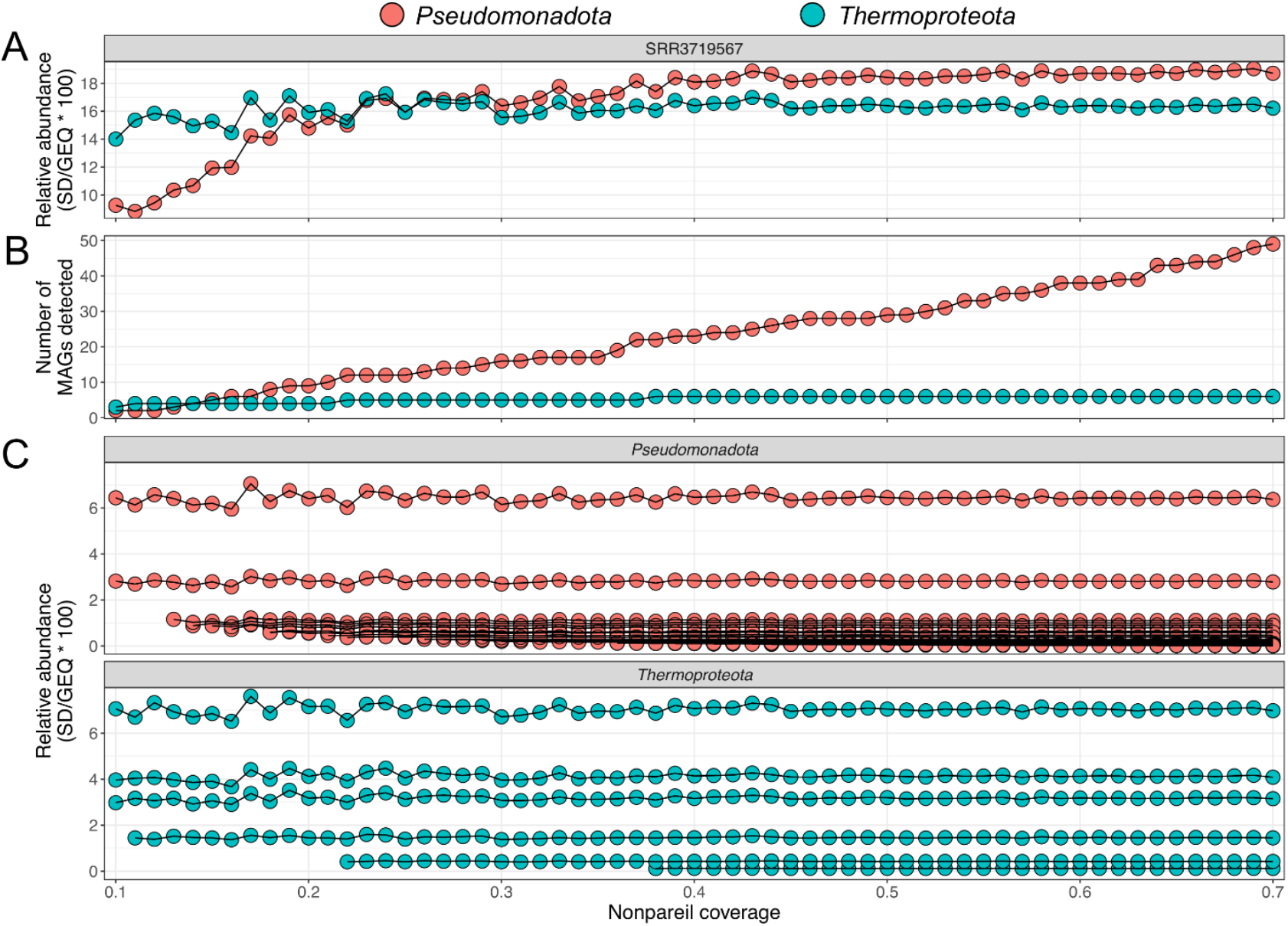
Effect of Nonpareil coverage on the comparison of relative abundance of two phyla based on a single metagenome. A) Changes in aggregated relative abundance (y-axis) of MAGs belonging to *Pseudomonadota* and *Thermoproteota* phyla as a function of Nonpareil coverage (x-axis) of subsampled datasets of the metagenome (SRR3719567) provided by Hawley and colleagues. Note that while the abundance of *Pseudomonadota* seems to be overall higher than that of *Thermoproteota* at high coverage datasets, *Thermoproteota* appear to be more abundant (i.e., the reverse result was obtained) in subsampled datasets of similar low coverage (e.g., Npc <0.2) due to several low abundance (not detectable at this level) *Pseudomonadota* MAGs (see panels B and C). B) Number of MAGs of each phylum detected (y-axis) as a function of Nonpareil coverage (x-axis). Note the continuous increase in the number of *Pseudomonadota* MAGs detected as Nonpareil coverage increases (and the corresponding increase in their aggregated relative abundance in A) while the number of *Thermoproteota* MAGs detected remains much more stable in the same datasets. These results are attributed to *Thermoproteota* encompassing only a few but relatively abundant MAGs that are detectable in low coverage datasets (hence, their aggregated relative abundance is reliably calculated even in the latter datasets) whereas *Pseudomonadota* includes many low abundant MAGs that are undetected, especially in low-to-medium coverage subsamples (hence, they do not contribute to the phylum’s aggregated abundance). C) Relative abundance (y-axis) of individual MAGs (each line represents one MAG) belonging to each phylum (colors) as a function of Nonpareil coverage (x-axis). Note that the relative abundance of individual MAGs remains stable across sub-samplings, supporting that abundance is reliable at any Npc as long as the MAG is detectable.

Given that the relative abundance of a group of MAGs (or genes) comprising a feature can vary with coverage (e.g., Npc), the outcomes of differential abundance analyses may also vary accordingly, providing for spurious or unreliable results. This bias can be minimized by subsampling (normalizing) to the same Npc, analogously to rarefaction. To exemplify the latter with more complex metagenome comparisons than those mentioned above, we focused on the vertical depth profile of *Marinisomatales* in the same data by Hawley and colleagues (5). A set of subsampled metagenomes with an even Npc produced a peak in abundance at 150 meters, in agreement with the full highly covered metagenomes, while the profile based on uneven Npc revealed the opposite trend as an effect of coverage, not actual biological/ecological differences (**Figure S1**). We also observed that subsampled metagenomes with even coverage at medium-to-high Npc (>=0.5) showed similar depth-profile trends compared to the full metagenomes (average Npc=0.8) while those at 0.4 Npc or below displayed divergent trends (**Figure S2**). This observation is in line with the 0.6 Npc threshold previously proposed to ensure adequate coverage of the sampled community and thus, meaningful biological comparisons between metagenomes (4). Similarly for features that involve genes, Zhang *et al*., 2021 (8) apparently overestimated the antibiotic resistance gene (ARG) diversity found in effluent water samples of wastewater treatment plants (WWTP) due to the higher Npc of these samples compared to influent and sludge samples (**Figure S3AB**). Consistently, half of the ARGs subtypes identified as differentially abundant between different WWTPs displayed a significantly different pattern when the metagenomes were subsampled to the same Npc (**Figure S3AC**). Such cases are probably widespread in recent microbiome literature. Based on a literature search, the number of manuscripts performing differential abundance analysis with metagenomic data has increased from 89 manuscripts in 2014 to 2,880 in 2023 (**Figure S4**). However, only 11 manuscripts out of the total 2,880 (0.38%) considered Nonpareil in their analyses. In addition, popular statistical tools for differential feature abundance analysis of amplicon data, such as metagenomeSeq and edgeR (9, 10), apply different approaches to normalize read counts based on library size (e.g., CSS and TMM) but do not incorporate differences in gene/genome length nor coverage making them unsuitable for metagenomic data.

We next opted to answer the question of how similar, in terms of Npc, the metagenomes should be in order to provide reliable comparisons. For this, we calculated the maximum difference in Npc, as a fraction of the sample’s Npc, that provided statistically identical results between the original and the subsampled metagenomes for a feature (ΔNpc_max_; see Methods); i.e., unbiased results. The histogram of ΔNpc_max_ values showed a wide and even distribution with no predominant peaks (**Figure S5**), suggesting that this parameter is highly variable and there is no universal cutoff. Instead, we observed that the ΔNpc_max_ positively correlated with the average relative abundance of the feature of interest for various metagenomes from different environments (R^2^ 0.57, p-value <0.001, **Figure S6**), revealing no sample- or habitat-specific biases. These findings suggested that assessing abundant features, and thus robustly detected even in subsampled metagenomes, may not be biased when comparing metagenomes of different Npc while low abundance features are more likely to be biased even when the ΔNpc is relatively low. For example, the estimate of abundance for a taxon with an abundance of 0.1% (measured as SD/GEQ*100) of the total metagenome will not be biased unless ΔNpc is higher than 50%, based on the marine metagenomes with medium-to-high Npc used here. In contrast, the estimate for a taxon with an abundance of 0.02% will be biased if the ΔNpc is higher than 10%. The correlations shown on **Figure S6** can be useful to identify situations where this bias will be important to take into account or not.

Sequencing high-coverage metagenomes (e.g., Npc 0.9-1) is the most effective way to minimize coverage-associated biases. However, obtaining high coverage is not practical for several environments, such as soils, or studies with many samples. For such cases, data normalization is likely necessary for comparative purposes, but determining the optimal normalization approach in any given situation may be challenging. To assist researchers with this challenge, we provide a decision tree that can be used to guide comparative metagenomic analyses under different scenarios (**Figure 2**). Specifically, researchers need to be cautious when the metagenome coverage is not high (e.g., Npc <0.9) and there are differences between the Nonpareil coverage of the metagenomes being compared. In such cases, we recommend first calculating the ΔNpc_max_ of the feature of interest (a single MAG/gene or a group of MAGs/genes representing a taxon/function) based on the metagenome(s) with the highest Npc in the comparison performed using the Npc_max.R script (see data availability statement). If the ΔNpc_max_ is lower than the actual ΔNpc between the metagenomes compared, the comparison might be biased due to differences in coverage. In the latter case, we recommend calculating the relative abundance of the feature in metagenomes that have been normalized by subsampling to the same Npc (e.g., to the lowest Npc) (**Figure 2**). Relative abundance and/or diversity values obtained after Npc normalization can then be used for more reliable comparative analyses. Metagenome subsampling and read mapping to calculate abundances in subsampled metagenome can be computational expensive and requires experienced users. To assist with this step, we provide a fast, easy-to-use script (Npc_normalizer.R) that accurately estimates relative abundance in subsampled metagenomes at a given, user-defined Npc using the abundance in the original complete metagenome. The abundance predictions of our tool showed perfect correlation (R^2^=1) with minimal error (∼1%) when compared to the observed abundance values obtained by actually subsampling and mapping the metagenomes reads (**Figure S7**), and thus can be used for direct comparisons between metagenomes.

**Figure 2.**
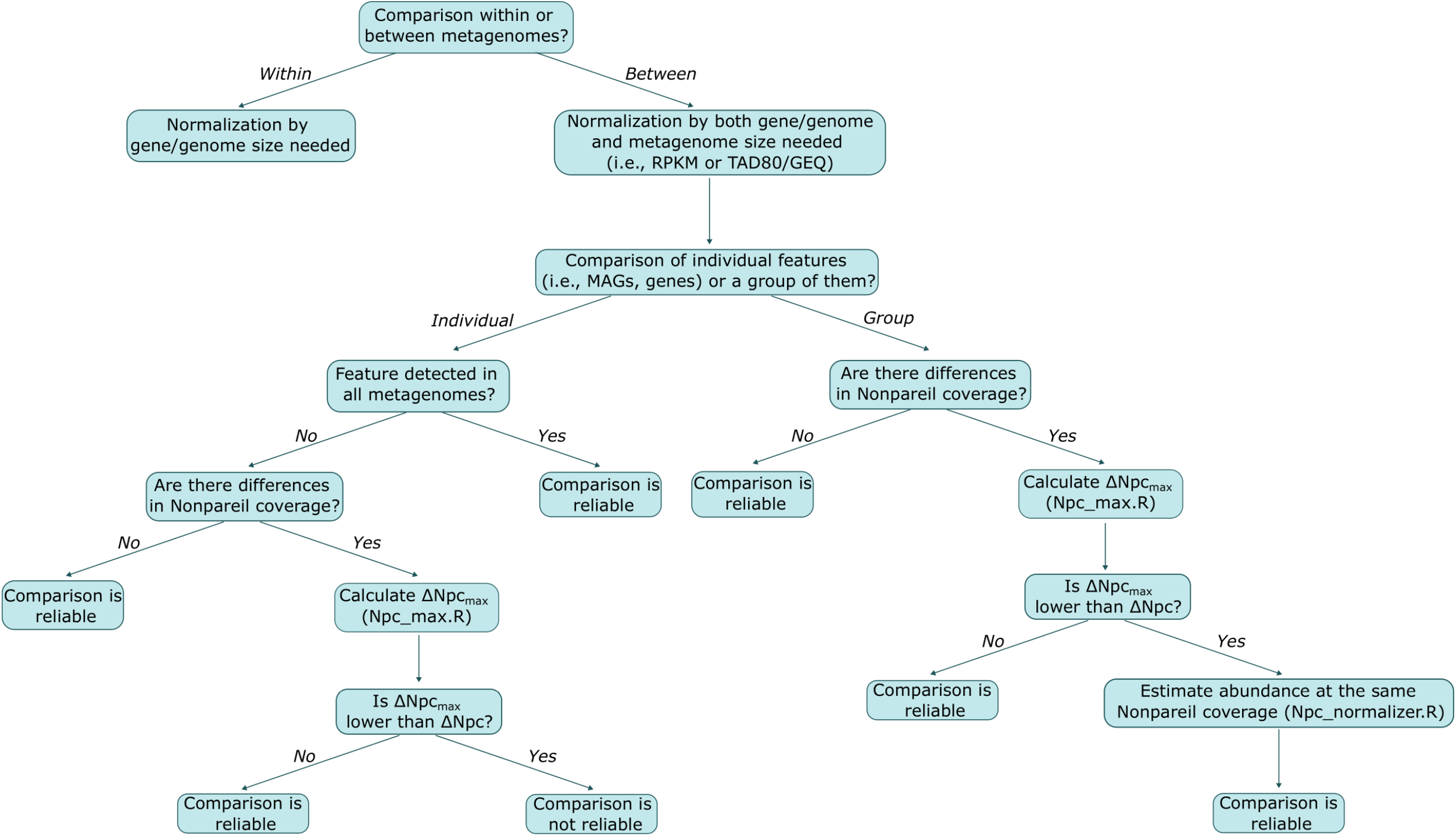
Decision tree to guide analysis of differential feature abundance or diversity between metagenomes.

It should be noted that even after Npc coverage normalization (for metagenomes of uneven coverage) the relative abundances of features obtained could differ somewhat from the actual abundances in the completely sequenced samples as exemplified by the results shown in **Figure 1**. Thus, coverage normalization can make comparative analyses more robust but cannot produce absolute estimates, especially if normalizing to relatively low coverage (e.g., Npc<0.2; see **Figure 1A**), and the level of its success depends on the abundance distribution of the members of the feature being assessed. The abundance distribution is typically unknown; therefore, dealing, in full, with this limitation is currently not feasible or requires more data (e.g. Npc coverage >0.9). Further, despite some controversy regarding the potential loss of statistical power with data subsampling (1, 11), avoiding false positives commonly outweighs the risk of missing significant comparisons (false negatives), making rarefaction a pragmatic and advantageous approach compared to the alternative of analyzing the data as is. Moreover, since the abundance predictions generated by the Npc_normalizer.R script produce maximum-likelihood estimates based on the complete data, the expected central value for abundances is adjusted to be comparable across datasets without a corresponding increase in the estimation error. Indeed, we observed ∼2.1% average deviation within the observed sequencing depth values after 10 random subsampling of the same metagenome, which is in the same range or even higher than the ∼1% average deviation of the same abundance value estimated by our script. Therefore, we recommend using this coverage normalization approach and our method outlined above when possible, and resorting to random subsampling only for more complex cases not covered by our tool.

In summary, we suggest to: 1) aim to sequence high coverage metagenomes to minimize the fraction of the community not covered, 2) be aware of and calculate metagenome coverage, 3) normalize the data to the same Npc when there are substantial differences in Npc (e.g., ΔNpc_max_ < ΔNpc), and 4) be aware that results from differential abundance or diversity analyses for the obtained Npc may differ in higher coverage metagenomes. Accordingly, we expect that the recommendations provided will help to minimize biases in comparative metagenomic studies, thereby facilitating the generation of quantitative, standardized and meaningful results.

## Methods

### Metagenome quality filtering, assembly and binning

A total of 85 marine metagenomes representing surface to 200 meters deep previously published by Hawley and colleagues (5) were downloaded from NCBI using the prefetch and fastq-dump (--minReadLen 50 --qual-filter --split-spot) tools of the SRA toolkit. Raw reads were quality-filtered using bbduk.sh v38.18 (qtrim=w,3 trimq=17 minlength=70 tbo tossjunk=t cardinalityout=t; https://jgi.doe.gov/data-and-tools/software-tools/bbtools/). Reformat.sh v38.18 separated pair-end reads in two files and Nonpareil v3.4.1(3) was employed to calculate metagenome coverage and diversity on cleaned forward reads with the options -T kmer -f fastq -X 50000 -t 8. Cleaned reads were assembled with SPAdes v3.15.5 (--meta --only-assembler -t 24 -k 21,33,55,77,99,127) (12) and contigs longer than 1kb were selected for binning with MaxBin2 v2.2.7 (13) and metaBAT2 v2.15 (14). Metagenome-assembled genomes (MAGs) were dereplicated with dRep v3.4.3 (-comp 50 - con 15 -sa 0.95) (15) yielding a total of 219 species-like MAGs that were quality-assessed using CheckM v1.2.2 (16) and taxonomically classified with GTDB-tk v2.3.2 (r214) (17).

### MAG relative abundance

Relative abundance was calculated as Sequencing Depth (SD) divided by Genome Equivalents (GEQ), which normalizes for differences in the sequencing effort applied, length of MAGs/targets, and average genome size between the metagenomes(18) as well as for spurious matches(19, 20). For this purpose, SD and GEQ were estimated by CoverM v0.6.1 (genome -p bwa-mem --min-read-percent-identity 95 --min-read-aligned-percent 70 --min-covered-fraction 10 --exclude-supplementary -m mean --output-format sparse -t 12)(21) and MicrobeCensus v1.1.0(18) (default settings), respectively. CoverM was also used to calculate mapped read counts which were converted to relative abundance as Read Per Kilobase per Million reads (RPKM). For simplicity, we used “relative abundance” to refer to the SD/GEQ metric, unless noted otherwise; e.g., for RPKM data. Plots were drawn in R using the ggplot2 library (22).

### Nonpareil coverage normalization

Metagenomes were subsampled to Nonpareil coverage (Npc) values ranging from 0.1 to 0.7 in 0.01 increments. For this, the number of reads needed to reach a given Npc was calculated with the predict.Nonpareil.Curve function from the Enveomics R library. Then, reads were randomly subsampled to the target number of reads using reformat.sh. The approach was implemented in the Npc_normalizer_manual.R script. Users can use this script to get randomly subsampled metagenomes with a given Npc and subsequently perform read mapping to features, etc. Alternatively, to simplify and speed up Npc normalization and reduce estimation error, relative abundance or count values can also be estimated using the Npc_normalizer.R script. The .npo files from the Nonpareil analysis, the original relative abundance, GEQ or count values and the MAG/gene length should be provided to run the script. The proposed normalization approach consists of the following steps:

1. Estimation of the fraction of reads required to achieve a given Npc for each metagenome. For this purpose, the predict.Nonpareil.Curve function is used to calculate the number of reads at the target Npc, which is then divided by the total number of reads in the original metagenome.
2. The fraction obtained is multiplied by sequencing depth, GEQ or reads counts in the original metagenome to get the estimated relative abundance values in the subsampled metagenomes.
3. Finally, only MAGs/genes with a minimum 0.1X sequencing depth, which equates to ∼10% breadth coverage according to the Lander-Waterman equation, are considered as present and used for comparisons between metagenomes. The remaining MAGs/genes are considered as undetected (i.e., zeros are assigned to MAGs/genes with sequencing depth <0.1X).

### Maximum acceptable difference in Npc (ΔNpc_max_)

To calculate the ΔNpc_max_ in a systematic fashion, we used the t.test function from stats R package v4.2.0 to compare the relative abundance at 0.7 Npc (reference) of each taxon at the phylum, class, order, and family level against the relative abundance of the same taxon in subsampled metagenomes showing stepwise decreasing Npc values (0.01 Npc per step). We ran this loop until the t.test became statistically significant. The difference between 0.7 and the specific Npc of the subsample considered was termed ΔNpc_max_, which was divided by the reference Npc (0.7) to calculate the maximum percentage of difference in Npc for statistically significant differences in feature abundance (shown on **Figure S5 and S6**). For features with only one member, ΔNpc_max_ was calculated based on the minimum Npc at which that member was detected. This analysis was performed using the seawater metagenomes described above as well as the human gut, freshwater and peat soil metagenomes published in Kim *et al*., 2022 (23), Rodriguez-R *et al*., 2020 (20) and Duchesneau *et al*., 2024 (24), respectively. The approach was implemented in the Npc_max.R script, which can be used to calculate the ΔNpc_max_ of custom user data.

### Antibiotic resistance genes relative abundance in wastewater metagenomes

To showcase the impact of Npc normalization in avoiding biased biological results due to differences in coverage, the dataset analyzed in Zhang *et al*., 2021 (8) was reanalyzed here following the same pipeline they used but introducing normalization. Briefly, raw reads were downloaded with prefetch and fastq-dump and cleaned with fastp v0.21.0 (default parameters). Nonpareil coverage was calculated, and metagenomes subsampled to the same Npc. The abundance of antibiotic resistance genes in the original as well as the subsampled metagenomes was calculated with ARGs-OAP v3.2.4 (25) (default parameters) as also performed in the study by Zhang and colleagues. The estimateR function from the vegan R package (26) was employed to calculate Chao1 index using as input the unnormalized counts for ARG subtypes provided by ARGs-OAP. Differential abundance test was performed with the t_test function of the R package rstatix (27) using the ARG subtypes abundance normalized by cell counts. Plots were drawn on R with ggplot2 v3.4.2 (22).

## Data availability

The scripts developed for this manuscript, along with instructions about how to run them, are available at https://github.com/baldeguer-riquelme/Npc_normalizer.

## Acknowledgments

This work has been supported, in part, by the US Department of Energy (Award No DE-SC0023297) to KTK.

