## Supplementary material for "Differences in metagenome coverage may confound abundance-based and diversity conclusions": SI appendix

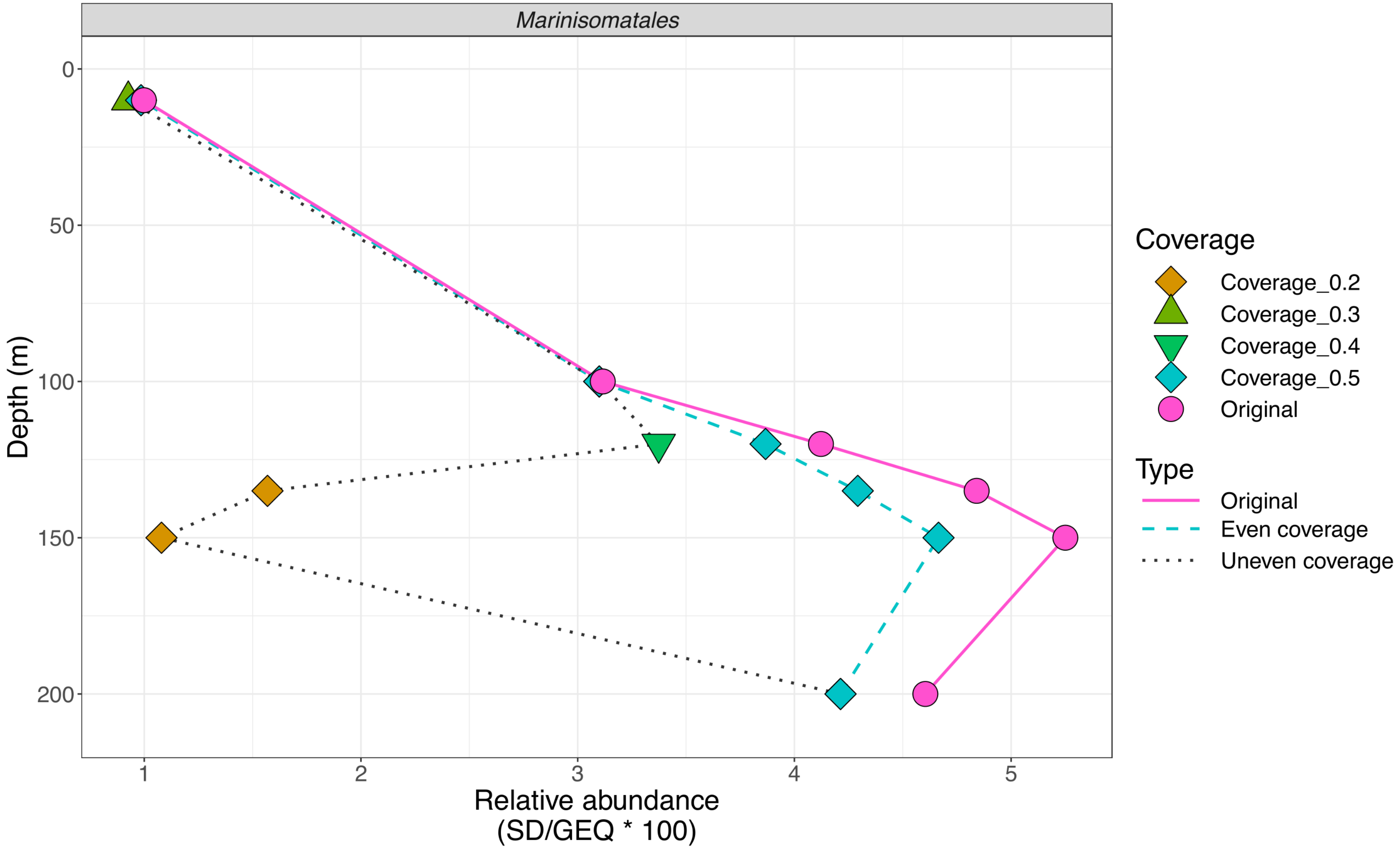


**Figure S1. Comparison of abundance trends across depth in the original (full-size) metagenomes (solid line, average Npc=0.8) and subsampled metagenomes at the same (dashed line) and different Nonpareil (dotted line) coverage levels.** The plot represents the aggregated relative abundance (x-axis) of MAGs belonging to the order *Marinisomatales* along the depth profile of the ocean (y-axis) based on the metagenomes provided by Hawley and colleagues. Note the substantially different trends in the even vs. the uneven coverage metagenomes.

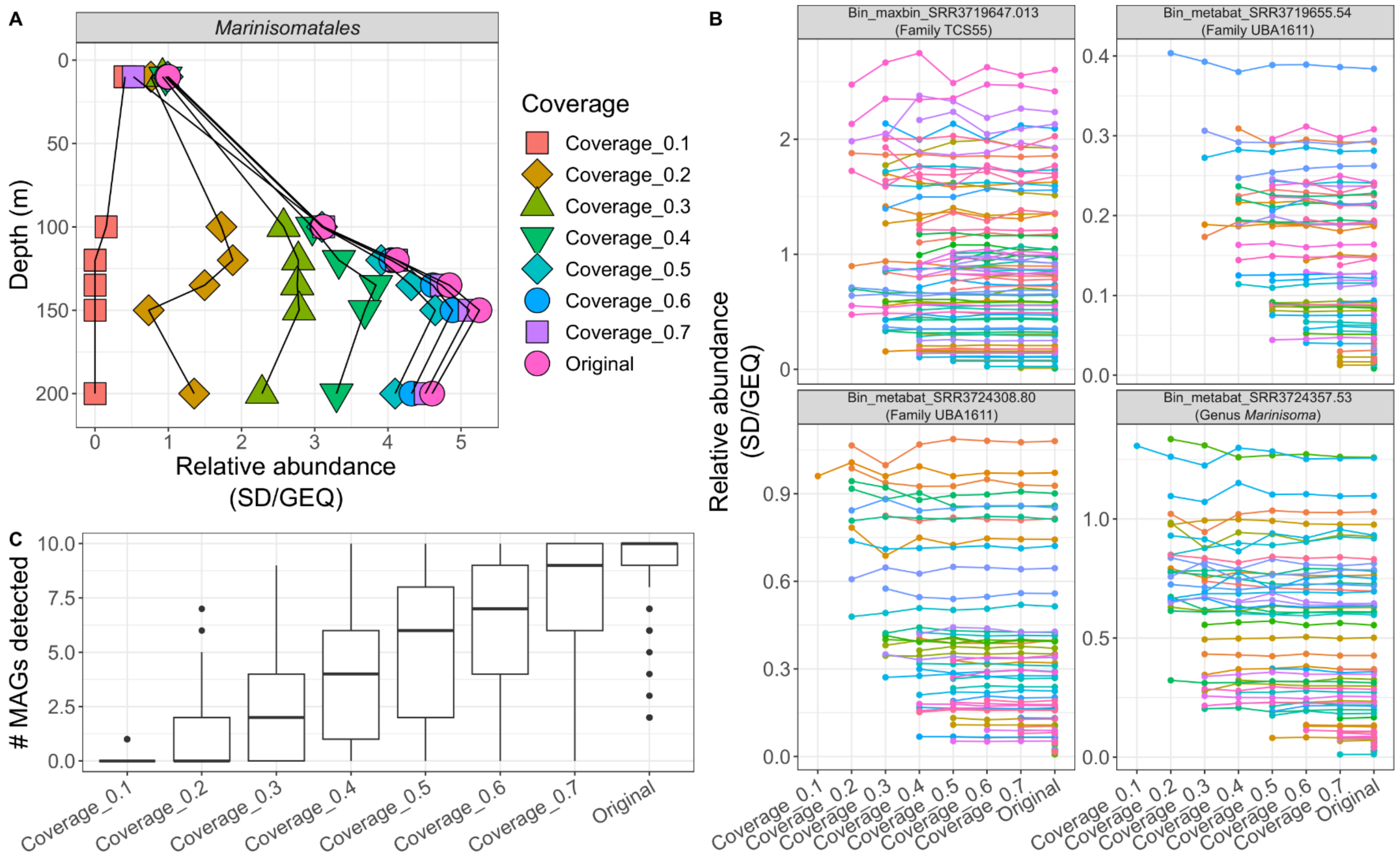


**Figure S2. Effect of metagenome coverage on the aggregated relative abundance of a feature.** A) Aggregated relative abundance (x-axis), calculated as sequencing depth (SD) divided by genome equivalents (GEQ), of a group of MAGs belonging to the order *Marinisomatales* is shown (x-axis) against the depth of the water column that the corresponding metagenomes were obtained from (y-axis). The average abundances from replicate metagenomes at each depth (10m n=8 replicates; 100m n=15; 120m n=12; 135m n=11; 150m n=15; 200m n=16) and subsampled metagenomes are shown. Note the increase in relative abundance as metagenome coverage increases. B) Relative abundance (y-axis) of individual MAGs belonging to *Marinisomatales* across subsampled metagenomes (x-axis). Each panel represents an individual MAG and each line one sample. Note the consistency of relative abundance values across subsampled metagenomes but also that several MAGs become undetectable at low coverage subsamples (and thus, after this point, they do not contribute to the relative abundance of the *Marinisomatales* order). C) Number of *Marinisomatales* MAGs (y-axis) detected in subsampled metagenomes (x-axis). Note the increase in the number of MAGs detected as metagenome coverage increases.

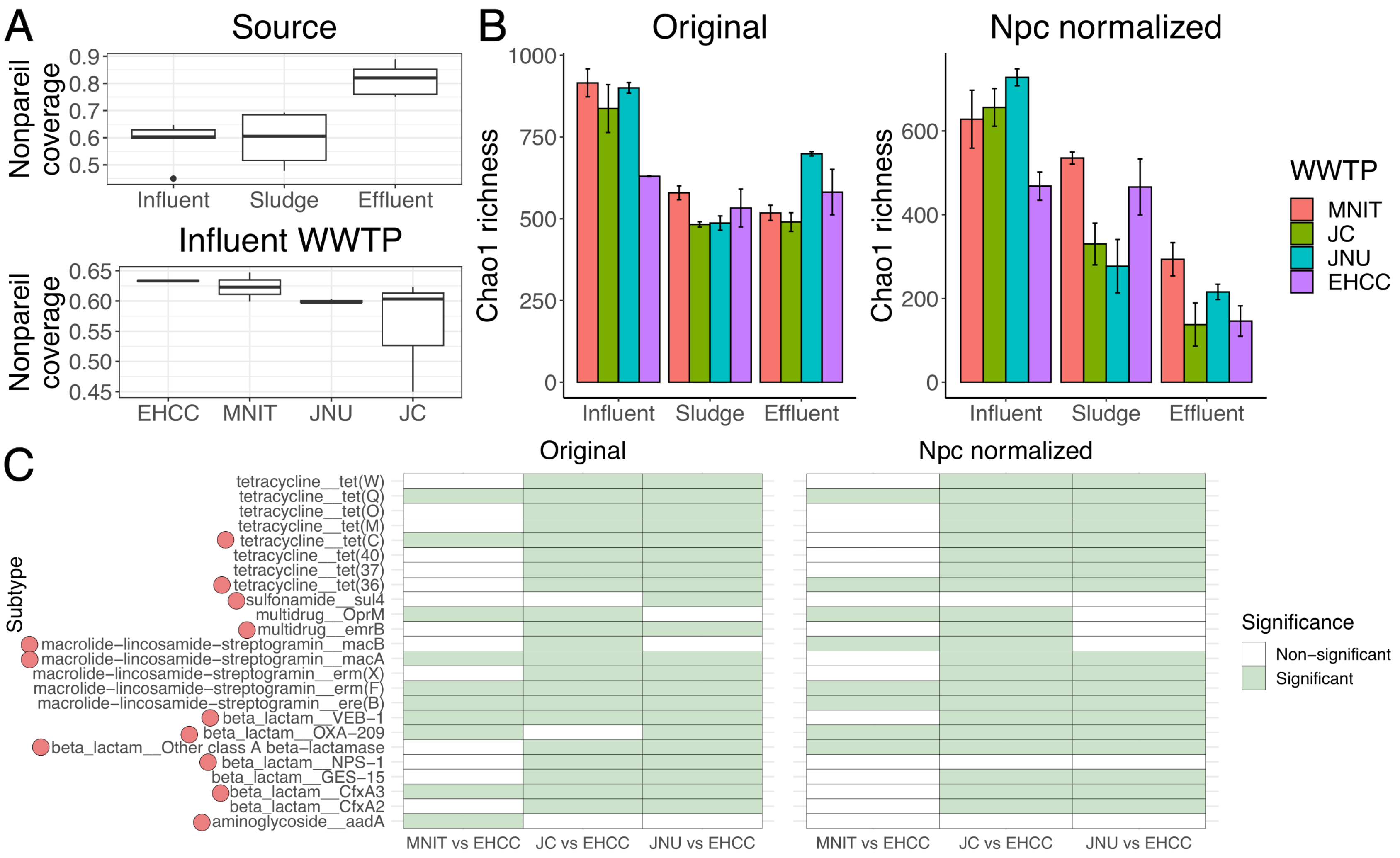


**Figure S3. Reanalysis of the data by Zhang and colleagues (2021) showcases the effect of uneven coverage on inferring differential abundance of antibiotic resistance gene (ARG) subtypes.** A) Nonpareil coverage (y-axis) of wastewater metagenomes collected from various stages (influent, sludge and effluent; x-axis top panel) of four different wastewater treatment plants (WWTP; x-axis bottom panel). Note the higher Nonpareil coverage of effluent metagenomes compared to influent and sludge metagenomes as well as the different nonpareil coverage for each WWTP. B) Chao1 richness (y-axis) of ARGs in influent, sludge and effluent metagenomes (x-axis) of the four

WWTP (colors) calculated for the original metagenomes and the Nonpareil coverage normalized metagenomes (Npc=0.45, the lowest Npc in the dataset). Error bars show the standard deviation of the 3 replicates per source and WWTP (influents from MNIT and EHCC had 2 replicates). Note that ARGs richness between sludge and effluents was similar in the original metagenomes but effluents showed significantly lower richness after correcting by Nonpareil coverage. Richness of ARGs in the effluent samples was thus overestimated by the original study due to higher coverage of the latter samples (see A). C) Differential abundance analysis of ARGs subtypes (y-axis) in the original and the Nonpareil coverage normalized metagenomes obtained from influent samples. The EHCC WWTP was compared to the three other WWTP (MNIT, JC and JNU). Green cells indicate statistically significant differences (p-value < 0.05) using the Welch's test while white cells indicate no statistically significant differences. The subtypes shown here correspond to the same subtypes that were shown as statistically significant in the original publication (some ARG subtypes could not be analyzed here due to the different version of the ARG-OAP database used). Note that the results with the original and the Nonpareil coverage normalized metagenomes were different for about half of the ARG subtypes (see dots next to the ARG subtype name), indicating that uneven coverage could often lead to misleading results.

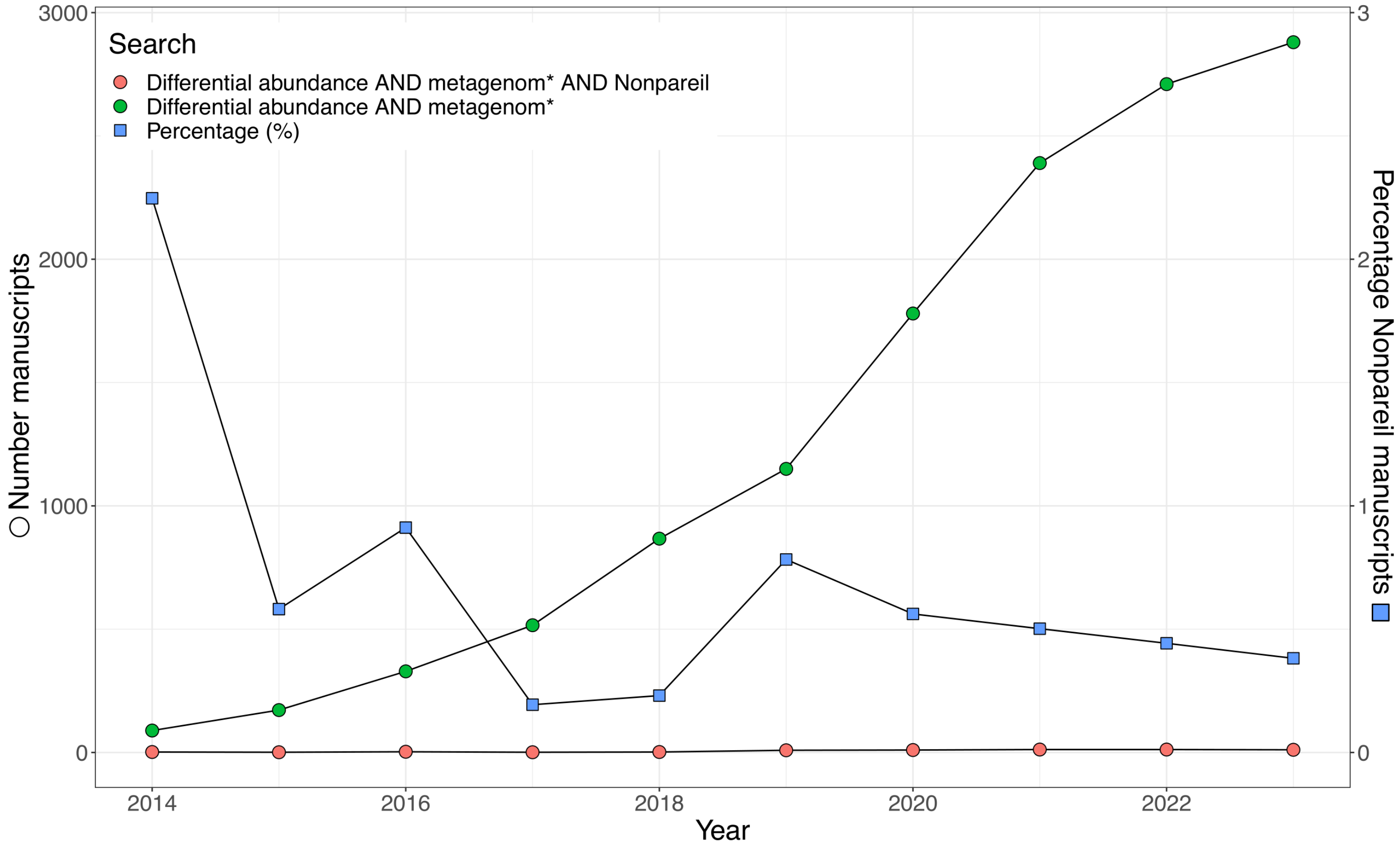


**Figure S4. Number of manuscripts (y-axis) published in the last 10 years (x-axis) that mentioned metagenomic differential abundance analysis, including how many also mentioned the Nonpareil tool.** The search was performed on Google Scholar on August 8th 2024, using the strings shown in the legend. Dots represent the number of manuscripts that, in addition to differential abundance and metagenomics, also mentioned (red) or not (green) Nonpareil. These values were used to calculate the percentage of manuscripts (blue squares; secondary y-axis) that mentioned Nonpareil from the total number of manuscripts that mentioned metagenomic differential abundance analysis. Note that Nonpareil was not mentioned in more than 99% of the manuscripts.

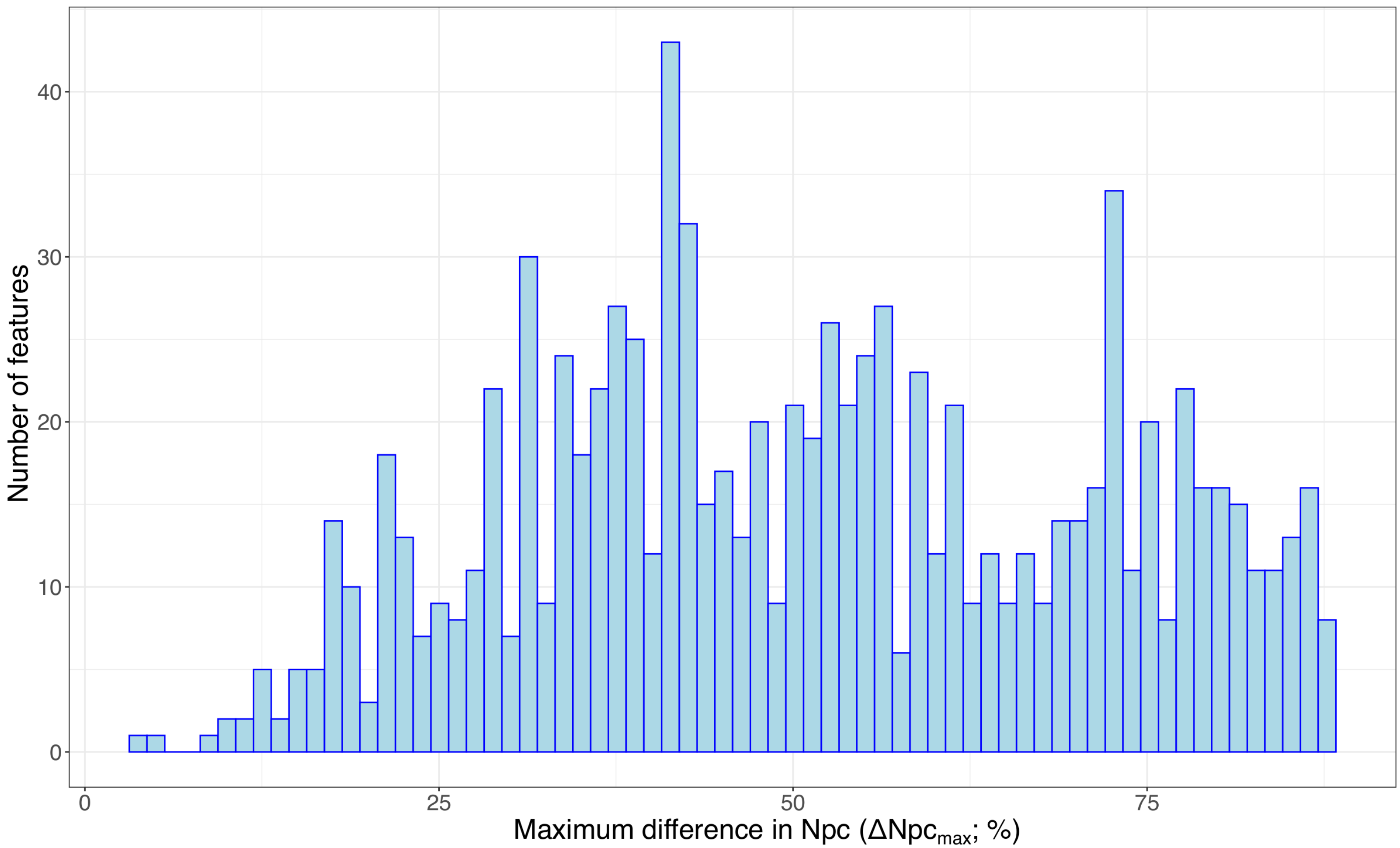


**Figure S5. Histogram of the maximum difference in Nonpareil coverage ($\Delta Npc_{max}$) between subsampled metagenomes that provided unbiased estimations of relative abundance differences compared to the full metagenomes.** A feature represents a group of MAGs belonging to the same taxonomic group: Phylum, Class, Order or Family). The graph shows the number of features (y-axis) that provided unbiased abundance results in subsampled metagenome relative to the full metagenome plotted against their corresponding $\Delta Npc_{max}$ value (x-axis) based on the marine metagenomes provided by Hawley and colleagues are displayed (n=1,289 features). Note the wide distribution obtained with not a clear threshold in $\Delta Npc_{max}$. See the Methods section for details of the analysis performed to calculate $\Delta Npc_{max}$ and main text for additional description of the results obtained.

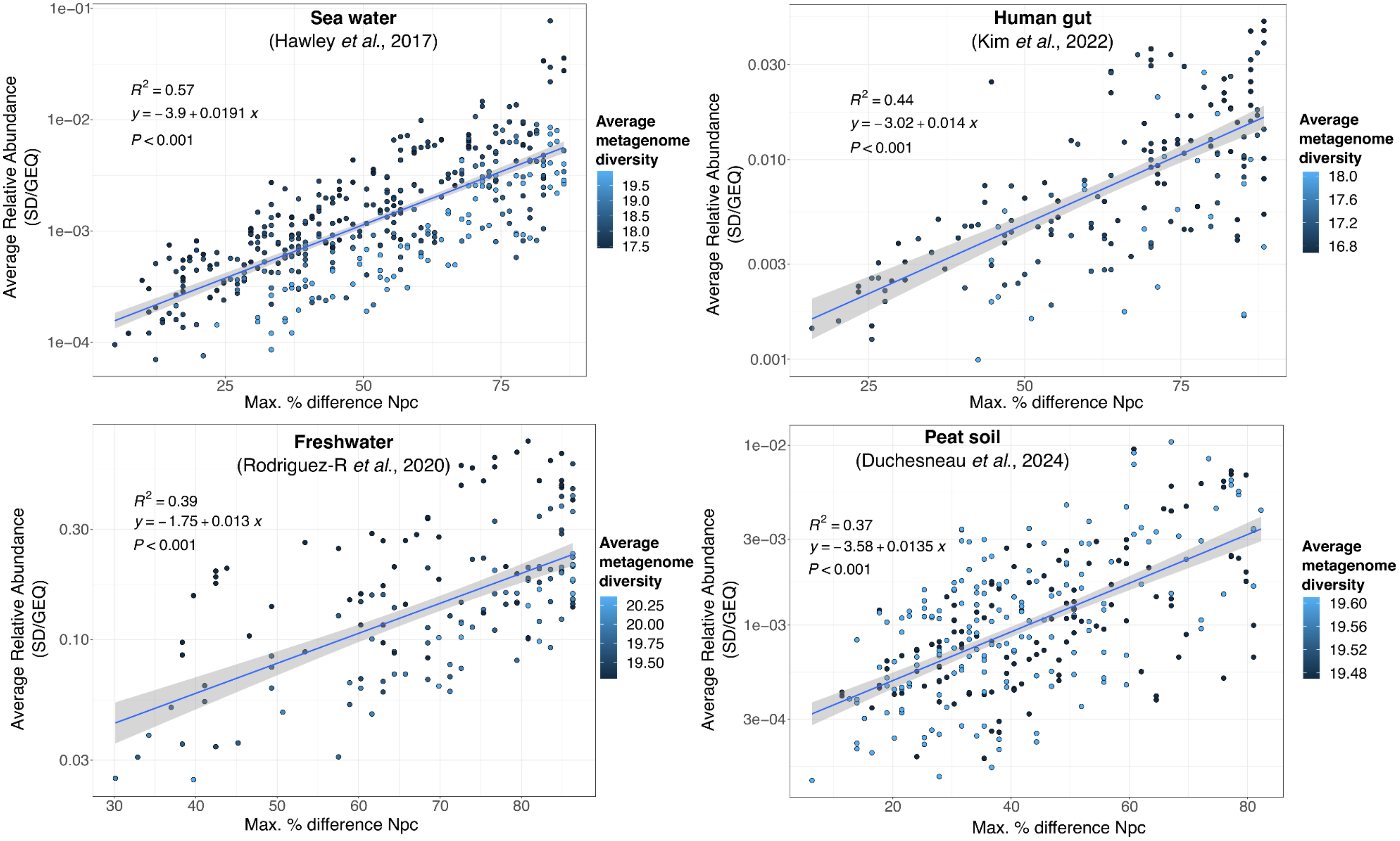


**Figure S6. Relationship between average relative abundance (x-axis) and maximum Npc difference for unbiased results (y-axis).** Each dot represents one feature, which represents a group of MAGs belonging to the same taxonomic group (Phylum, Class, Order or Family). The dot represents the average aggregated abundance of the members of the feature (x-axis) in a group of metagenomes (e.g., metagenomes from the same depth in A or same lake in C) relative to the maximum acceptable difference in Npc ($\Delta Npc_{max}$ in %; y-axis) to obtain the same abundance estimate as in the original metagenomes (meaning statistically unsignificant by Welch's test). $\Delta Npc_{max}$ is expressed as a fraction (percentage) of the Npc value of the original, non-subsampled metagenome. The analysis was performed with four different environments: seawater (A), human gut (B), freshwater (C) and peat soils (D) to cover a wide range of habitats and Nonpareil diversity values (from 16.8 to 20.25). Colors in dots represent the average Nonpareil diversity for the group of metagenomes used in the analysis. Note that the positive correlation was strong and consistent for all four environments. The equation provided in each plot can be used to calculate the maximum acceptable difference in Npc (in %) for performing abundance comparison between two metagenomes (or their subsamples) based on a given relative abundance value of a target feature.

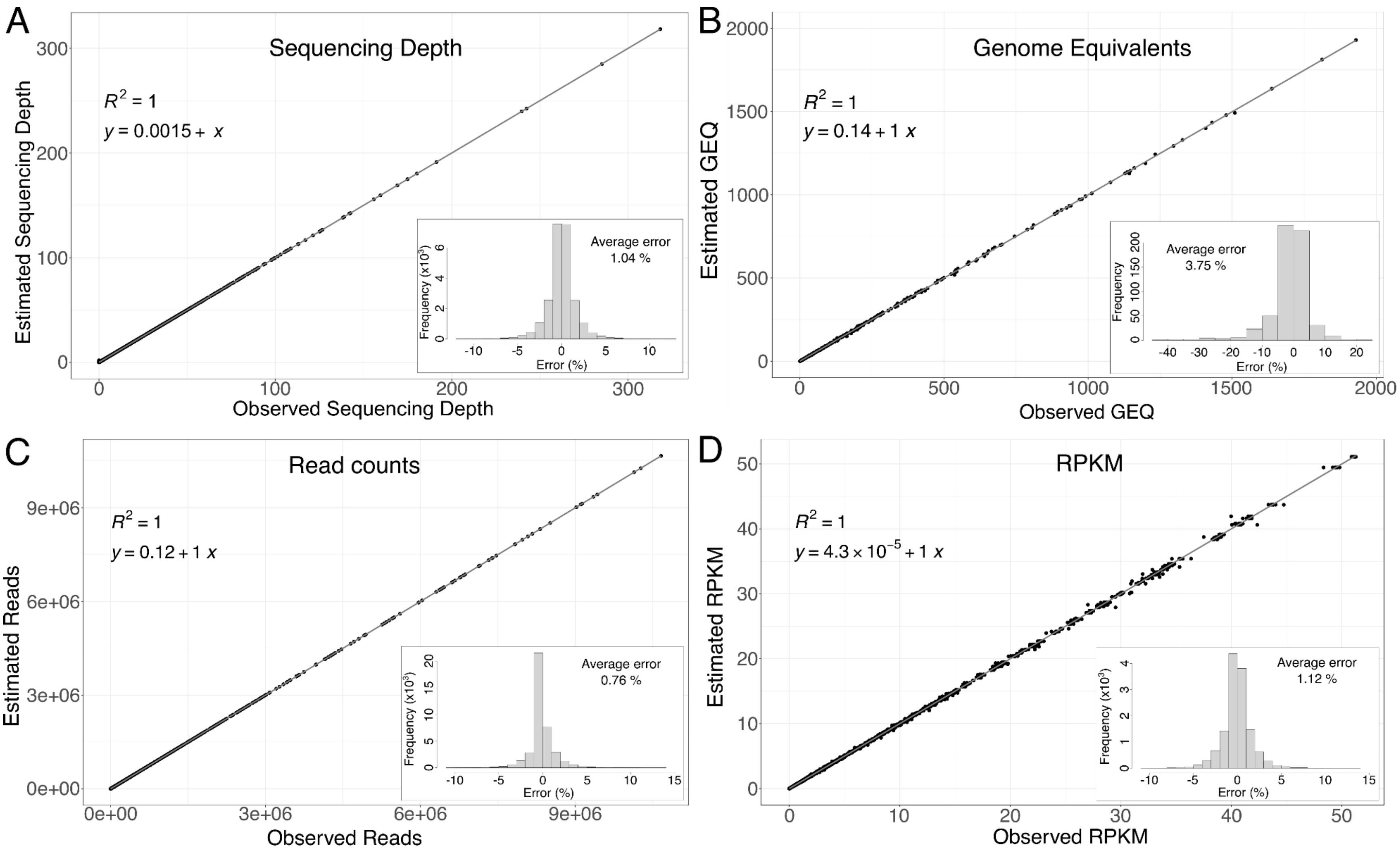


**Figure S7. Accuracy of the relative abundance and read count estimation in subsampled metagenomes obtained with the Npc_normalizer.R script.** The accuracy was assessed for several metrics, namely Sequencing Depth (panel A), Genome Equivalents (GEQ; panel B), read counts (panel C) and RPKM (panel D). For each metric, dots represent the estimated value of each (group of) MAG(s) (panels A, C and D) or metagenome (for GEQ in panel B) calculated by the script (y-axis) compared to the actual observed value (x-axis) obtained by manually subsampling and mapping the subsampled metagenomes to the same reference MAG dataset. The estimated values for the four metrics obtained by the script show an almost perfect correlation against the observed values ($R^2$=1) and very low percentages of error (~1%; bar plots at the top left of each panel). The marine metagenomes obtained by Hawley and colleagues was used for these tests.
